# The genetic architecture of human cortical folding

**DOI:** 10.1101/2021.01.13.426555

**Authors:** Dennis van der Meer, Tobias Kaufmann, Alexey A. Shadrin, Carolina Makowski, Oleksandr Frei, Daniel Roelfs, Jennifer Monereo Sánchez, David E.J. Linden, Jaroslav Rokicki, Christiaan de Leeuw, Wesley K. Thompson, Robert Loughnan, Chun Chieh Fan, Paul M. Thompson, Lars T. Westlye, Ole A. Andreassen, Anders M. Dale

## Abstract

The folding of the human cerebral cortex is a highly genetically regulated process that allows for a much larger surface area to fit into the cranial vault and optimizes functional organization. Sulcal depth is a robust, yet understudied measure of localized folding, previously associated with a range of neurodevelopmental disorders. Here, we report the first genome-wide association study of sulcal depth. Through the Multivariate Omnibus Statistical Test (MOSTest) applied to vertexwise measures from 33,748 participants of the UK Biobank (mean age 64.3 years, 52.0% female) we identified 856 genetic loci associated with sulcal depth at genome-wide significance (α=5×10^-8^). Comparison with two other measures of cortical morphology, cortical thickness and surface area, indicated that sulcal depth has higher yield in terms of loci discovered, higher heritability and higher effective sample size. There was a large amount of genetic overlap between the three traits, with gene-based analyses indicating strong associations with neurodevelopmental processes. Our findings demonstrate sulcal depth is a promising MRI phenotype that may enhance our understanding of human cortical morphology.

---

During early brain development, the cerebral cortical sheet folds into gyri and sulci in a highly regulated manner, due to multiple intrinsic and extrinsic mechanical forces.^1–3^ This cortical folding not only allows for a much larger surface area to fit into the cranial vault, but also reduces distance between neurons, leading to faster signal transmission.^2^ Accordingly, measures of sulcal morphology are associated with cognitive performance^4^ and lack of cortical folding (lissencephaly) is accompanied by severe mental retardation.^5^ Atypical folding can result from defects in neuronal proliferation, migration, and differentiation, and has been associated with major neurodevelopmental^6–8^ and neurodegenerative disorders.^9^

Sulcal depth is a rather understudied measure of sulcal morphology, reflecting the convexity or concavity of any given point on the cortical surface. This measure is very robust, and captures complex localized folding patterns of the cerebral surface.^10^

Several studies have indicated there is a strong genetic component to sulcal depth, which is mostly prenatally determined.^11,12^ Sulci are more similar in monozygotic than dizygotic twins,^13^ and an estimated 56% of between-subject variance in average depth of the central sulcus is under genetic control.^14^ Further, Williams syndrome, caused by deletion of a section of chromosome 7, is associated with widespread reductions in sulcal depth,^15^ which mediate its behavioral symptoms.^16,17^ Yet, there has been no large-scale molecular genetics study of this measure.

Here, we provide the first genome-wide association study (GWAS) of sulcal depth, comparing its genetic architecture to the more commonly studied brain morphological measures of cortical thickness and surface area. Given that gene variants are likely to have distributed effects across MRI phenotypes, we targeted a multivariate analysis of a vertex-wise representation of the cortical surface. To that end, we performed the Multivariate Omnibus Statistical Test (MOSTest)^18^ on data from 1153 vertices, using a common template (fsaverage3), with the medial wall vertices excluded. Our sample consisted of 33,748 unrelated White European participants of the UK Biobank (UKB), with a mean age of 64.3 years (standard deviation (SD) 7.5 years), 52.0% female.

Following surface reconstruction, we pre-residualized all vertices for age, sex, scanner site, a proxy of image quality (FreeSurfer’s Euler number),^19^ and the first twenty genetic principal components to control for population stratification. After applying a rank-based inverse normal transformation, MOSTest was performed on the resulting residualized measures, yielding a multivariate association with 9.1 million included SNPs (see online methods for more details). We additionally repeated the main GWAS analyses while covarying for the mean across all vertices in order to remove global effects; the findings from these analyses were highly similar to the main analyses, as reported in the supplements.

## Discovery

MOSTest revealed 856 independent loci reaching the genome-wide significance threshold of a=5×10^-8^ for sulcal depth, see Figure 1a. In comparison, for surface area and thickness we found 661 and 591 loci, respectively, see Supplementary Figure 1. The effects of discovered top variants followed gyral and sulcal patterns, as shown in Figure 1b for the most significant SNP at chromosome 15. Data S1-S3 contains information on all discovered loci per trait, including mapped genes, and lists the significance of each lead SNP for the other two traits as well.

**Figure 1.**
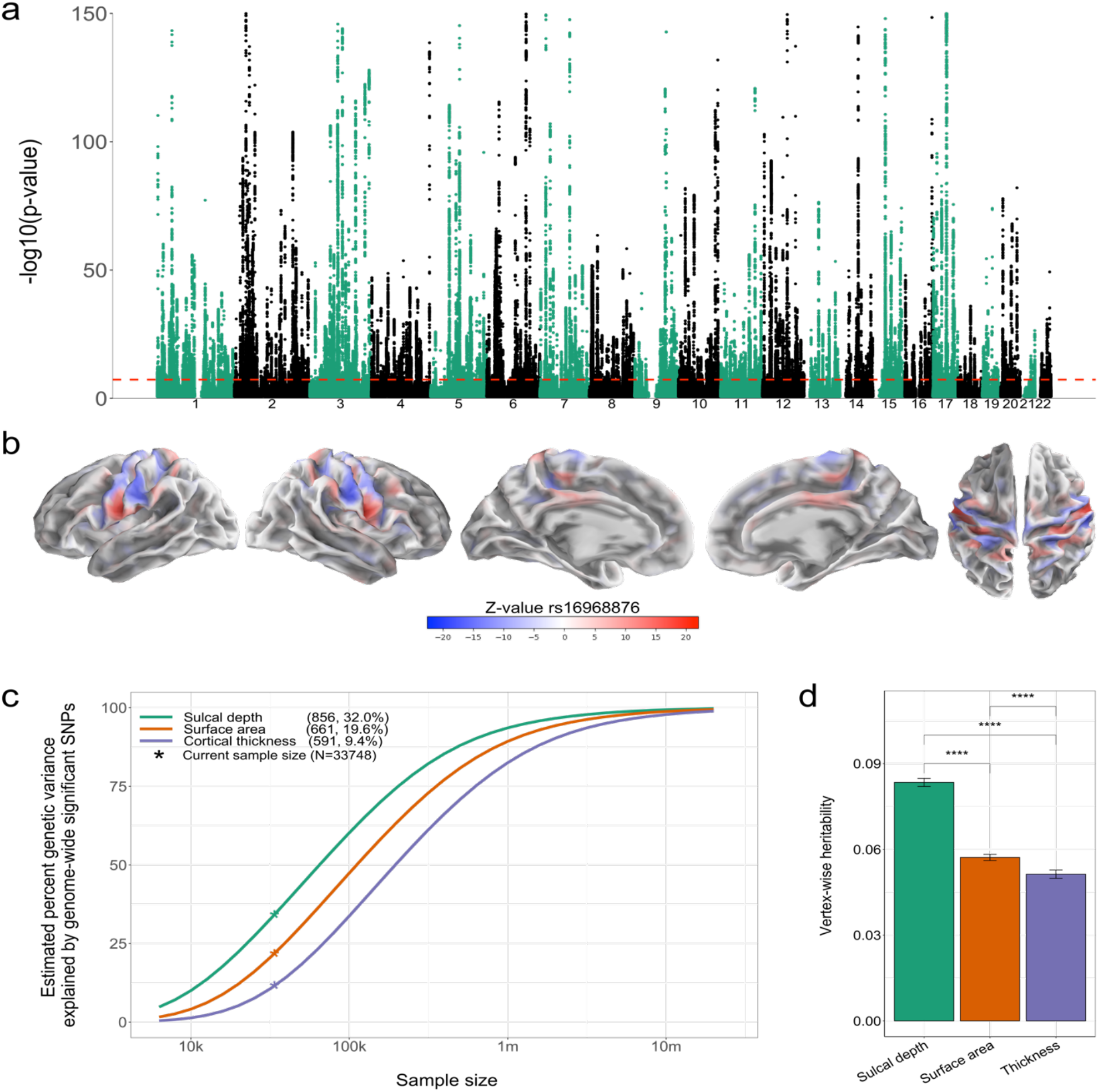
Locus discovery. **a)** Manhattan plot of the multivariate GWAS on sulcal depth, with the observed - log10(p-value) of each SNP shown on the y-axis. The x-axis shows the relative genomic location, grouped by chromosome, and the red dashed line indicate the whole-genome significance threshold of 5×10^-8^. The y-axis is clipped at −log10(p-value)=150. **b)** Brain map depicting the vertex-wise z-values for the top hit rs16968876 at chromosome 15. **c)** Power plot, showing the relation between variance explained by genome-wide significant hits (y-axis) and sample size (x-axis). The number of hits discovered per modality and the percent explained genetic variance is indicated between brackets in the legend. **d)** Bar plot of the mean SNP-based heritability (with 95% confidence interval) across vertices (on the y-axis) per modality (x-axis). In c) and d), sulcal depth is represented in green, surface area in orange and cortical thickness in purple.

Next, using the MiXeR tool,^20,21^ we fitted a Gaussian mixture model of the null and non-null effects to the three GWAS summary statistics, estimating the polygenicity and effect size variance (‘discoverability’). The results are summarized in Figure 1c, depicting the estimated proportion of genetic variance explained by discovered SNPs for each trait as a function of sample size. The horizontal shift of the curve across the different traits indicates that the effective sample size is the highest for sulcal depth and lowest for cortical thickness. Further, the mean heritability of sulcal depth, over all the vertices, was significantly higher than for the two other traits, see Figure 1d, i.e. the higher genetic signal in sulcal depth is also captured by univariate measures.

## Genetic overlap

Next, we analyzed the genetic overlap between the three traits at the locus level, gene level and pathway level. At the locus level, we found that 625 loci had overlapping start and end positions between sulcal depth and surface area (Dice coefficient of .82), 509 loci overlapped between sulcal depth and cortical thickness (Dice=.70), and 450 loci overlapped between surface area and thickness (Dice=.72). 447 loci overlapped across all three traits, see Figure 2a.

**Figure 2.**
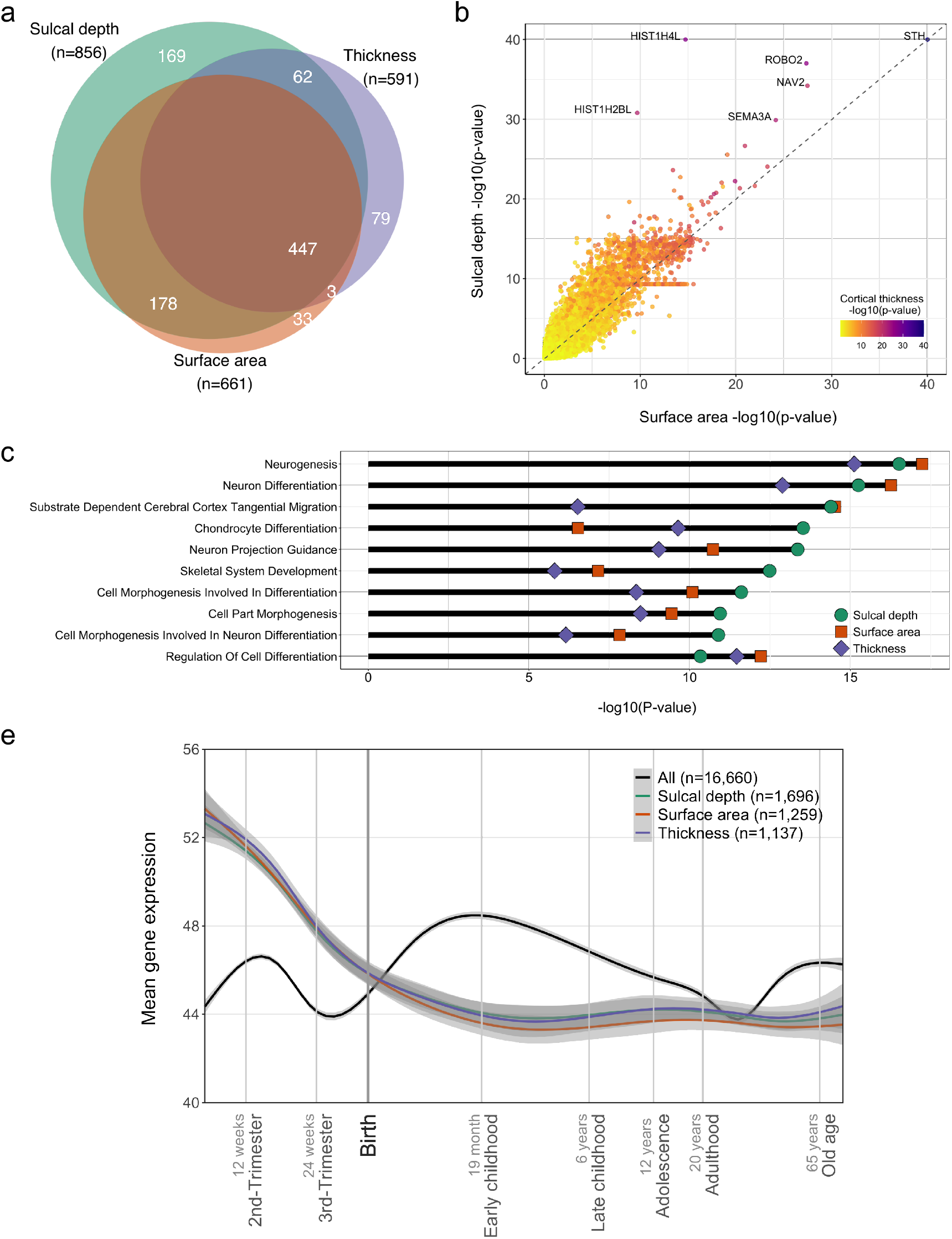
Genetic overlap. **a)** Venn diagram of the amount of discovered loci overlapping between the three different traits. **b)** Scatterplot of gene-based p-values, with y-axis indicating p-values for sulcal depth, x-axis those for surface area and the coloring indicating p-values for cortical thickness. Note, −log10(p-values) are clipped at 40. **c)** Ten most significant gene pathways for sulcal depth, as listed on the y-axis, with the −log10(p-values) indicated on the x-axis for each of the three traits. **d)** Mean, normalized expression (y-axis) of genes over time (x-axis, log10 scale) per trait and over all available genes, as indicated by colors. Grey shading indicates 95% confidence bands.

The large genetic overlap between the traits was also evident at the gene level, as illustrated in Figure 2b. The top gene *STH*, thought to play a role in phosphorylation of tau,^22^ was highly significantly associated with all three traits. *ROBO2*, *NAV2* and *SEMA3A*, key players in neuronal outgrowth guidance,^23–25^ were also associated with all three traits. The two histone genes *HIST1H4L* and *HIST1H2BL* were relatively specifically associated with sulcal depth; histone is central in regulating brain development through its role in gene expression.^26^

Figure 2c shows the top ten most significant Gene Ontology pathways for sulcal depth, together with its p-values for the two other traits. We found strong associations with neurogenesis and neuron differentiation pathways, overlapping between all three traits. Associations with neuronal tangential migration were shared by sulcal depth and surface area but much less by cortical thickness, in line with the role of tangential migration of neurons in determining cortical folding.^27^ Notably, pathways related to chondrocyte differentiation and skeletal system development appeared more specific to sulcal depth, possibly pointing towards early life interactions between cortical folding and the shaping of the cranium.^2^

We further coupled the findings of our genebased analyses to cortical gene expression patterns, derived from post-mortem brain tissue of clinically unremarkable donors across the lifespan.^28^ As shown in Figure 2d, the probes tagging genes associated with the three traits showed a distinct profile over the lifespan, characterized by high prenatal expression and low postnatal expression.

## Discussion

Here, we reported the results from the first large-scale molecular genetics study of sulcal depth. With 856 loci discovered, explaining an estimated 32% of its genetic variance, this study has found the highest number of loci for any brain trait considered so far.

The direct comparison with surface area and thickness indicated that sulcal depth is more heritable and its genetic determinants are more discoverable. This may reflect the evolutionary significance of cortical folding, the development of which enabled the advent of a larger brain and optimization of its functional organization.^29^ A synthesis of the literature suggests that human-specific folding follows from an interplay between mechanical forces and cellular mechanisms that have come about over the course of evolution through mutations of genes primarily coupled to cell cycling and neurogenesis.^30^ Our findings suggest that the sulcal depth metric is closely aligned with these genetic processes that shape highly important brain morphological characteristics.

As indicated by the brain maps, genetic effects may have opposing directions of effects on neighbouring points in the brain and are not likely to be captured by ROIs as defined in common parcellation schemes. This is in line with strong differences in the morphology and arrangement of neurons and fibres along cortical folds, varying widely from the gyral crown along the lateral wall down to the sulcal fundus.^30,31^ This speaks to the use of vertex-wise data to maximally capture such patterns. The presence of widespread, complex genetic effects also attests to the application of multivariate tools to boost discovery of genetic determinants by leveraging shared signal between measures.

We further found large genetic overlap between all three morphological brain traits, extending our previous findings that surface area and thickness share the majority of their genetic determinants.^18,32^ This is in contrast to other studies which suggested that surface area and thickness are genetically independent of each other.^33,34^ Those studies made use of genetic correlation estimates, while the approach used here does not rely on consistent directions of effects across the genome, which is unlikely to be the case for pairs of complex traits.^21,35^ We found that sulcal depth overlaps more with surface area than with cortical thickness, indicating a closer relation between the degree of cortical folding and surface area. However, these metrics do partly capture distinct genetic processes, i.e. sulcal depth is likely to provide additional information on the genetics of brain morphology to complement what is found through studies of surface area and cortical thickness.

In addition to the reported locus overlap, the specific identified genetic variants, genes and pathways further inform our understanding of cortical morphology and associated disorders. The most significant pathways were particularly relevant for early brain development, with neurogenesis and differentiation ranking highest. This fits very well with a large body of literature on the genetic regulation of the mechanical forces that drive cortical folding.^30^ It is also in accordance with our findings that the sets of identified genes showed highest expression in fetal cortical tissue. Further, cortical folding has been shown to take place almost entirely prenatally,^12^ with sulcal patterns at birth being predictive of neurobehavioral outcomes.^11^ Follow-ups on our work with neuroimaging data across the lifespan, including infants, is needed to replicate these findings and to further determine spatiotemporal patterns of genetic effects on sulcal depth. Given reported associations of sulcal morphology with a range of neurodevelopmental and neurodegenerative disorders, ^6–9^ it will also be of interest to investigate how sulcal depth genetics relates to the development of brain disorders.

To conclude, despite the evolutionary and ontogenetic importance of cortical folding, sulcal depth is an underexplored trait that is genetically more discoverable than cortical thickness and surface area. Further investigation of this trait may significantly enhance our understanding of the human brain and associated disorders.

## Supporting information

DataS1-6

## Author contributions

D.v.d.M, O.A.A. and A.M.D. conceived the study; D.v.d.M., A.A.S and T.K. pre-processed the data. D.v.d.M. and A.A.S. performed all analyses, with conceptual input from A.M.D., D.R., J.R., C.d.L., O.A.A., L.T.W. and T.K.; All authors contributed to interpretation of results; D.v.d.M. drafted the manuscript and all authors contributed to and approved the final manuscript.

## Materials & Correspondence

The data incorporated in this work were gathered from public resources. The code is available via https://github.com/precimed (GPLv3 license). Correspondence and requests for materials should be addressed to

## Acknowledgements

The authors were funded by the Research Council of Norway (276082, 213837, 223273, 204966/F20, 229129, 249795/F20, 225989, 248778, 249795, 298646, 300767), the South-Eastern Norway Regional Health Authority (2013-123, 2014-097, 2015-073, 2016-064, 2017-004, 2019-101), Stiftelsen Kristian Gerhard Jebsen (SKGJ-Med-008), The European Research Council (ERC) under the European Union’s Horizon 2020 research and innovation programme (ERC Starting Grant, Grant agreement No. 802998), ERA-Net Cofund through the ERA PerMed project ‘IMPLEMENT’, and National Institutes of Health (R01MH100351, R01GM104400). This work was partly performed on the TSD (Tjeneste for Sensitive Data) facilities, owned by the University of Oslo, operated and developed by the TSD service group at the University of Oslo, IT-Department (USIT). Computations were also performed on resources provided by UNINETT Sigma2 - the National Infrastructure for High Performance Computing and Data Storage in Norway.

## Online Methods

### Participants

We made use of data from participants of the UKB population cohort, obtained from the data repository under accession number 27412. The composition, set-up, and data gathering protocols of the UKB have been extensively described elsewhere^36^. For this study, we selected White Europeans that had undergone the neuroimaging protocol. For the primary analysis, making use of T1 MRI scan data released up to March 2020, we excluded 771 individuals with bad structural scan quality as indicated by an age and sex-adjusted Euler number^19^ more than three standard deviations lower than the scanner site mean. We further excluded one of each pair of related individuals, as determined through genome wide complex trait analysis (GCTA), using a threshold of 0.0625 (n=1,138). Our sample size for this analysis was n=33,748, with a mean age of 64.3 years (SD=7.5). 52.0 % of the sample was female.

### Data preprocessing

T_1_-weighted scans were collected from three scanning sites throughout the United Kingdom, all on identically configured Siemens Skyra 3T scanners, with 32-channel receive head coils. The UKB core neuroimaging team has published extensive information on the applied scanning protocols and procedures, which we refer to for more details^37^. The T1 scans were obtained from the UKB data repositories and stored locally at the secure computing cluster of the University of Oslo. We applied the standard “recon-all −all” processing pipeline of Freesurfer v5.3., followed by extracting vertex-wise data for sulcal depth, surface area and thickness, at ico3 (1,284 vertices) and ico4 (5,124) resolution, without applying smoothing. We included both the left and right hemisphere measures and excluded non-cortical vertices, belonging to the medial wall.

Note that we have chosen sulcal depth as a metric of cortical folding, as this captures −vertex-wise-localized folding, providing the signed distance from the inflated surface.

We subsequently regressed out age, sex, scanner site, Euler number, and the first twenty genetic principal components from each vertex measure. Following this, we applied rank-based inverse normal transformation^38^ to the residuals of each measure, leading to normally distributed measures as input for the GWAS.

We reran the MOSTest analyses as described above, additionally regressing out the mean across all vertices for each of the three traits. The resulting number of loci are shown in Supplementary Table 1.

### MOSTest procedure

The MOSTest software is freely available at https://github.com/precimed/mostest, and details about the procedure and its extensive validation have been described previously.^18^ In brief, consider N variants and M (pre-residualized) phenotypes. Let z_ij_ be a z-score from the univariate association test between i^th^ variant and j^th^ (residualized) phenotype and z_i_ = (z_i1_,…, z_iM_) be the vector of z-scores of the i^th^ variant across M phenotypes. Let Z = {z_ij_} be the matrix of z-scores with variants in rows and phenotypes in columns. For each variant consider a random permutation of its genotypes and let 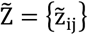 be the matrix of z-scores from the univariate association testing between variants with permuted genotypes and phenotypes. A random permutation of genotypes is done once for each variant and the resulting permuted genotype is tested for association with all phenotypes, therefore preserving correlation structure between phenotypes.

Let 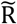 be the correlation matrix of 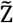, and 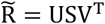 is its singular valued decomposition (U and V – orthogonal matrixes, S–diagonal matrix with singular values of 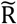 on the diagonal). Consider the regularized version of the correlation matrix 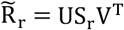, where S_r_ is obtained from S by keeping r largest singular values and replacing the remaining with r_th_ largest. The MOSTest statistics for i^th^ variant (scalar) is then estimated as 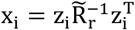, where regularization parameter r is selected separately for cortical area and thickness to maximize the yield of genome-wide significant loci. In this study we observed the largest yield for cortical surface area with r=10; the optimal choice for cortical thickness was r=20 (Supplementary Figure 5). The distribution of the test statistics under null 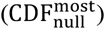 is approximated from the observed distribution of the test statistics with permuted genotypes, using the empirical distribution in the 99.99 percentile and Gamma distribution in the upper tail, where shape and scale parameters of Gamma distribution are fitted to the observed data. The p-value of the MOSTest test statistic for the i^th^ variant is then obtained as 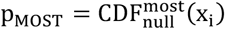.

### Univariate GWAS procedure

We made use of the UKB v3 imputed data, which has undergone extensive quality control procedures as described by the UKB genetics team^39^. After converting the BGEN format to PLINK binary format, we additionally carried out standard quality check procedures, including filtering out individuals with more than 10% missingness, SNPs with more than 5% missingness, and SNPs failing the Hardy-Weinberg equilibrium test at p=1*10^-9^. We further set a minor allele frequency threshold of 0.005, leaving 9,061,022 SNPs.

We have previously calculated that the number of features provided by fsaverage3, 1153 vertices following exclusion of the medial wall, leads to the maximum number of loci identified through MOSTest, compared to other resolutions.^40^ We followed up on the most significant findings from MOSTest through univariate GWAS on the 5124 vertices that make up fsaverage4, i.e. one level of resolution above fsaverage3. This was done to improve the resolution of the visualisations. The univariate GWAS on each of the pre-residualised and normalized measures were carried out using the standard additive model of linear association between genotype vector, g_j_, and phenotype vector, y. Independent significant SNPs and genomic loci were identified in accordance with the PGC locus definition, as also used in FUMA SNP2GENE.^41^ First, we select a subset of SNPs that pass genome-wide significance threshold 5×10^-8^, and use PLINK to perform a clumping procedure at LD r2=0.6, to identify the list of independent significant SNPs. Second, we clump the list of independent significant SNPs at LD r2=0.1 threshold to identify lead SNPs. Third, we query the reference panel for all candidate SNPs in LD r2 of 0.1 or higher with any lead SNPs. Further, for each lead SNP, it’s corresponding genomic loci is defined as a contiguous region of the lead SNPs’ chromosome, containing all candidate SNPs in r2=0.1 or higher LD with the lead SNP. Finally, adjacent genomic loci are merged together if they are separated by less than 250 KB. Allele LD correlations are computed from EUR population of the 1000 genomes Phase 3 data. We made use of the Functional Mapping and Annotation of GWAS (FUMA) online platform (https://fuma.ctglab.nl/) to map significant SNPs from the MOSTest analyses to genes.

We additionally performed clumping according to the definition used by the Enhancing NeuroImaging Genetics through Meta-Analysis (ENIGMA) consortium, to allow for comparison with previous imaging GWAS studies. According to this definition, loci were formed through PLINK using a p-value threshold of 5×10^-8^ (--clump-p1) and LD cutoffs of 1 Mb (--clump-kb) and r2< 0.2 (--clump-r2). Please see Supplementary Table 1 for the number of lead SNPs and loci according to both definitions.

Loci were defined to be overlapping between two traits if their start and end position overlapped. The Dice coefficient for each pair of traits was calculated as the number of overlapping loci divided by the sum of the total number of discovered loci for both traits.

### MiXeR analysis

We applied a causal mixture model^20,21^ to estimate the percentage of variance explained by genome-wide significant SNPs as a function of sample size. For each SNP, i, MiXeR models its additive genetic effect of allele substitution, β_i_, as a point-normal mixture, 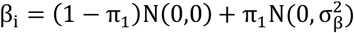, where π_1_ represents the proportion of non-null SNPs (`polygenicity`) and 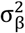 represents variance of effect sizes of non-null SNPs (`discoverability`). Then, for each SNP, j, MiXeR incorporates LD information and allele frequencies for 9,997,231 SNPs extracted from 1000 Genomes Phase3 data to estimate the expected probability distribution of the signed test statistic, 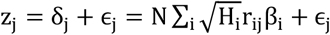, where N is sample size, H_i_ indicates heterozygosity of i-th SNP, r_ij_ indicates allelic correlation between i-th and j-th SNPs, and 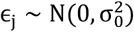 is the residual variance. Further, the three parameters, π_1_, 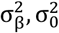, are fitted by direct maximization of the likelihood function. Fitting the univariate MiXeR model does not depend on the sign of z_j_, allowing us to calculate |z_j_| from MOSTest p-values. Finally, given the estimated parameters of the model, the power curve S(N) is then calculated from the posterior distribution p(δ_j_|z_j_, N).

### Gene-set analyses

We carried out gene-based analyses using MAGMA v1.08 with default settings, which entails the application of a SNP-wide mean model and use of the 1000 Genomes Phase 3 EUR reference panel. Gene-set analyses were done in a similar manner, restricting the sets under investigation to those that are part of the Gene Ontology biological processes subset (n=7522), as listed in the Molecular Signatures Database (MsigdB; c5.bp.v7.1).

Regarding the results from the gene-based analyses, in Figure 2, we note that there is a horizontal line visible at −p=5e-10, caused by many genes having this exact p-value. This is due to MAGMA switching to permutation when its numerical integration approach fails. MAGNA uses 1e-9 permutations, so when the observed is more extreme than this, this is the resulting p-value.

### Gene expression analyses

We made use of gene expression data derived from brain tissue from 56 clinically unremarkable donors ranging in age from 5 weeks post conception to 82 years.^28^ We took the data as preprocessed by Kang et al., selecting for each gene the probe with the highest differential stability, n=16,660. We subsequently averaged over 13 cortical regions, within donor, and normalized the expression values, within probe, across donors, to a range between 0 (lowest observed value) and 100 (highest observed value). Plotting of the mean expression over time per gene set was done with ggplot2 in R v4.0.3., with geom_smooth(method=“gam”) using default settings.

## Extended Data

### List of discovered loci and genes

Sup Please see Data S1-S3 for the full tables of discovered loci for each trait. This includes information on lead SNP, genomic location, significance, and mapped genes, as outputted by FUMA.

Data S4-S6 lists the genes found to be significant, following multiple-comparisons correction (α=.05/18,203), through MAGMA, per trait.

**Supplementary Figure 1.**
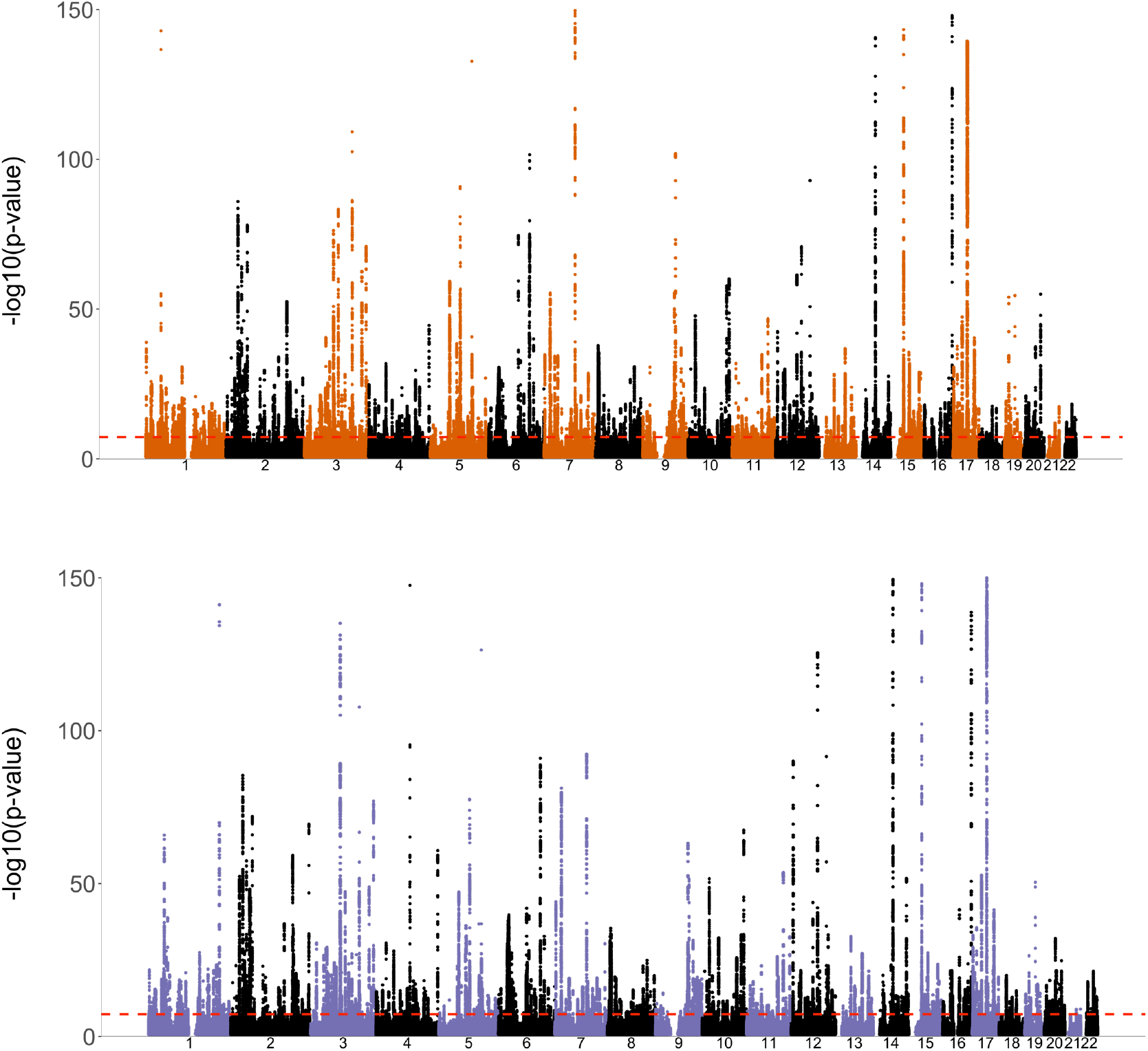
Manhattan plots for surface area (top, in orange) and cortical thickness (bottom, in purple). The observed −log10(p-value) of each SNP is shown on the y-axis. The x-axis shows the relative genomic location, grouped by chromosome, and the red dashed line indicate the whole-genome significance threshold of 5×10^-8^. The y-axis is clipped at - log10(p-value)=150.

**Supplementary Table 1.** Number of significant loci per trait for both PGC and ENIGMA locus definitions. Left: results from main analyses, right: results after additionally regressing out the mean across vertices (‘global’).

| Trait | Main analyses |  | Regressing out global |  |
| --- | --- | --- | --- | --- |
|  | PGC | ENIGMA | PGC | ENIGMA |
| Sulcal depth | 856 | 2242 | 880 | 2349 |
| Surface area | 661 | 1409 | 647 | 1369 |
| Thickness | 591 | 1173 | 580 | 1127 |

